# PeakClimber: A software tool for the accurate quantification of complex multianalyte HPLC chromatograms using the exponential Gaussian function

**DOI:** 10.1101/2024.08.05.606689

**Authors:** Joshua T. Derrick, Pragney Deme, Norman Haughey, Steven A. Farber, William B. Ludington

## Abstract

High-performance liquid chromatography (HPLC) is a common medium-throughput technique to quantify the components of often complex mixtures like those typically obtained from biological tissue extracts. However, analysis of HPLC data from complex multianalyte samples is hampered by a lack of tools to accurately determine the precise analyte quantities on a level of precision equivalent to mass-spectrometry approaches. To address this problem, we developed a tool we call PeakClimber, that uses a sum of exponential Gaussian functions to accurately deconvolve overlapping, multianalyte peaks in HPLC traces. Here we analytically show that HPLC peaks are well-fit by an exponential Gaussian function, that PeakClimber more accurately quantifies known peak areas than standard industry software for both HPLC and mass spectrometry applications, and that PeakClimber accurately quantifies differences in triglyceride abundances between colonized and germ-free fruit flies.

## Introduction

Liquid chromatography (LC) is a series of techniques to separate individual analytes from a mixture of chemicals using liquid phase solvent^1^. As compared to gas chromatography (GC), that uses inert gases as solvents (usually helium), liquid chromatography can separate particles of larger molecular weight^2^. Originally, liquid chromatography techniques relied on gravity for solvent flux, which meant that running of individual chromatographs took hours and sometimes days to complete. In the 1960s^3^, high-pressure (or performance) liquid chromatography (HPLC) was introduced, speeding up the flow rate by forcing the solvent through an extremely narrow column at high-pressures. Despite improvements in column performance, trade-offs between mass transfer resistance and diffusive behaviors fundamentally limit peak resolution^4^. For many HPLC applications, peak integration is sufficient, as these analyses principally are concerned with presence/absence of specific peaks or with quantitation of relatively pure analytes with little peak overlap. For the quantification of more complicated chemical and biological samples with overlapping peaks however, integration alone is inaccurate. Historically, this meant the operator spent considerable efforts to develop protocols to fully separate analyte peaks of interest, something not always possible.

Various solutions have been proposed to this problem. Common industry software, such as ThermoFisher’s Chromeleon and Water’s MassLynx utilize a method known as valley-to-valley,^5^ where a line is dropped from the lowest point between two peaks to the chromatograph’s baseline, which is determined by the rolling-ball method^6^. The two peaks are then integrated on either side of the line. This method has the advantages of being neutral to the underlying peak shape, independent of the surrounding peaks, and a fast runtime. However, most peaks map to some variation of the Gaussian distribution^4,7–12^ and are not independent of neighboring peaks with which they overlap. Two more recent open source software packages, HappyTools^13^ and hplc.io^14^, improve on the valley-to-valley method by fitting each chromatograph to a sum of Gaussian or skewed Gaussian curves, respectively. However, these theoretical peak shapes are not necessarily suited to the data, and the shape of a single peak is not universally agreed upon. Early quantitative models of liquid chromatography showed that analytes unbind the column with an exponential decay that is convolved by Gaussian noise based on their distribution along the length of the column and their diffusion in the liquid phase before reaching the detector^7,8,15–17^. While the shape of a peak depends on the amount of sample loaded on the column, Langmuir surface binding kinetics usually leads to a Gaussian shaped peak with tailing^18^.

In this manuscript, we show that HPLC analyte peaks are best fit with an exponential Gaussian function. Our tool, PeakClimber, fits chromatographs to a sum of exponential Gaussian curves. We show these curves are mathematically and empirically good fits for single analyte peaks and consistent with extensive literature suggesting that this approach empirically aligns with chromatography data^7,8,19–21^. PeakClimber also makes iterative improvements in denoising data, detrending data, and in reducing the runtime of the analysis. To highlight the utility of PeakClimber, we compare its performance to other algorithms by analyzing coinjections of three fatty-acids. Finally, PeakClimber was superior to traditional approaches in quantifying the differences in lipid composition between *Drosophila melanogaster* that were reared with and without bacteria.

## Results

### Traditional chromatography analysis methods fail to accurately quantify complex peaks

Valley-to-valley integration methods produce a mismatch between the calculated and true peak shape (Figure 1A). To quantify the error of this approach, we conducted a simulation with three-synthetic exponential Gaussian peaks with randomized parameters that overlapped significantly. Our simulations showed that the valley-to-valley method has significant error between the true peak shape and the valley-to-valley integration regions, but this error is especially marked for the first peak in the trace (Figure 1B). This is likely due to the undercounting of the exponential-tail region of the first peak by valley-to-valley analysis.

**Figure 1:**
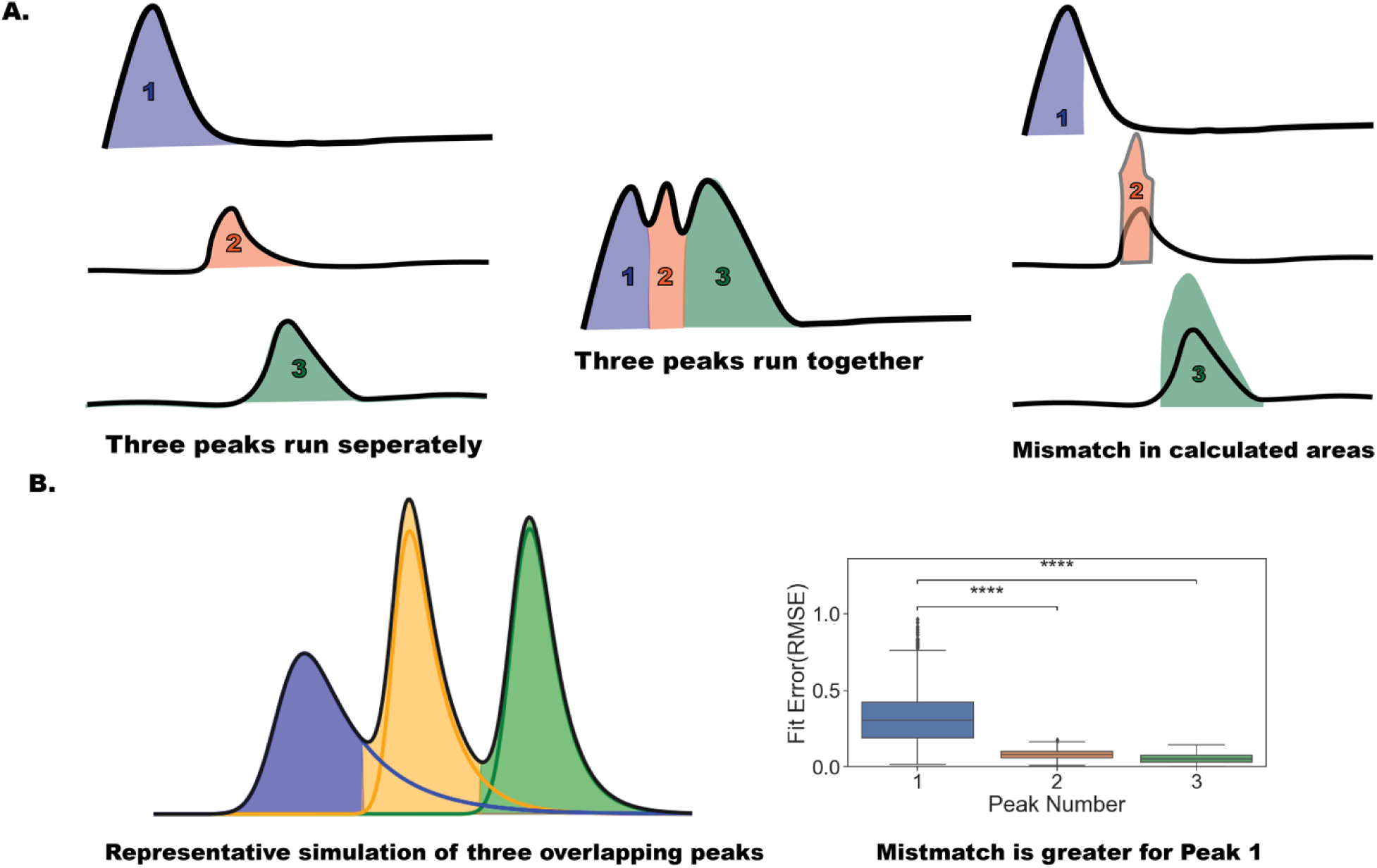
The problem of peak quantification. (**A**). A cartoon depiction of a common inaccuracy in peak quantification. When three analytes are well-separated, their area is accurately calculated by peak integration. When three analytes have overlap in the trace, the valley-to-valley area calculation algorithm will not accurately determine peak areas due to overlap of the tails. **(B)**. A simulation of three overlapping peaks. The shaded region represents the integration regions identified by the valley-to-valley algorithm; the solid lines represent the true peaks. The difference between the two is quantified by Root Mean Squared Error (RMSE) (n=1000, Mann-Whitney followed by Wilcoxon Ranked Test, ****p<1e-04).

### Single-analyte HPLC peaks fit an exponential Gaussian distribution

We first wanted to determine what shape we should use to fit individual peaks. There is extensive discussion of this question in the literature^1,4,5,8,8–12,15–19,22^, but there is broad agreement as to a generally Gaussian peak shape with some amount of tailing. To this end, we developed analytical, computational, and empirical models to support the exponential Gaussian as the true shape of a chromatographic peak.

#### Analytical and computational solutions

Consider a column of finite length, initially containing no solute. Injectant containing solute *S* is added to the column, and *S* is completely bound to the column at a single location. Solvent *U* is then run over the column. Solute *S* has affinity *k*_1_ for solvent *U*. We assume that the reverse reaction is negligible because unbound *S* flows away in the solvent. This behavior can be described by the differential equation:

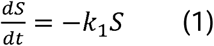

which we can solve analytically:

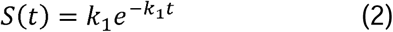

producing an exponential function. With an agent-based Monte-Carlo (Figure 2A, histogram) simulation with parameters for *S* (amount of analyte), *k*_1_ (affinity for solvent *U*), column length, and flow rate that are relevant to common HPLC columns, we recapitulate the analytical solution almost exactly (Figure 2A, red line on blue histogram).

**Figure 2:**
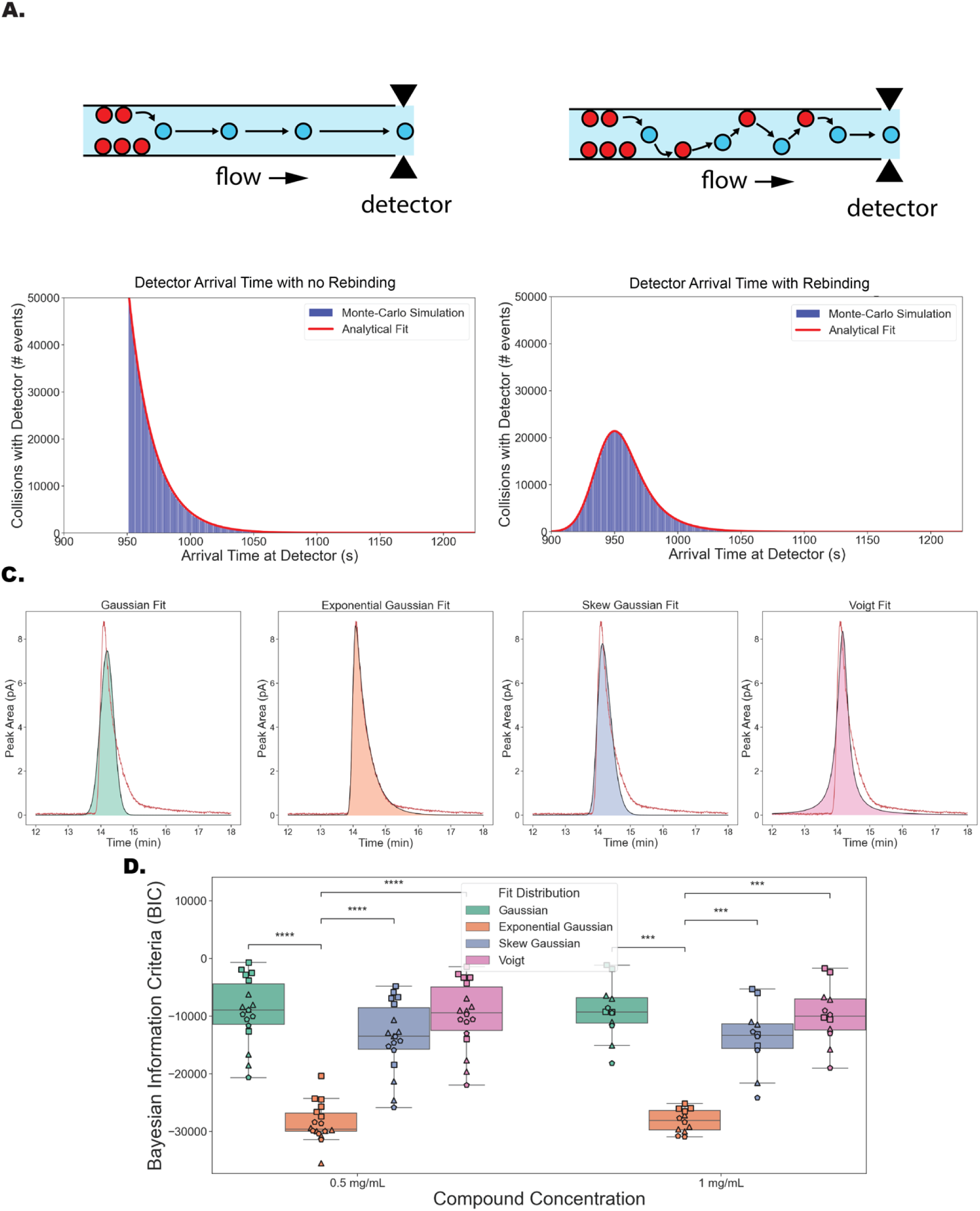
Individual HPLC analyte peaks correspond to an exponential Gaussian distribution. (**A and B**). Monte Carlo simulation of solute arrival time at the detector without (**A**) or with (**B**) rebinding (blue) fit to an exponential distribution (**A**) or exponential Gaussian distribution (**B**) (red) (n=10000 simulations consisting of 100 analyte molecules each). (**C**). Empirical fits of (left to right) Gaussian, exponential Gaussian, skew Gaussian, and Voigt distributions to a single linoleic acid peak (**D**). Bayesian Information Criteria (BIC) of the fit of above distributions on pooled arachidonic acid (triangle), docosahexaenoic acid (circle), and linoleic acid (square) single peaks. Analytes are grouped by injection volume. (n=12 [4 experimental replicates of each of the 3 fatty-acids], Mann-Whitney U-test with Bonferroni correction, ***: p<1e-03; ****: p<1e-04).

However, this initial model contains several incorrect assumptions, most notably that column binding and unbinding is a single event. In reality there are many binding and unbinding steps^10,17,23^. Thus, the distribution of analyte *S* will not be bound to a single site, but rather spread out across the column after many unbinding and binding events. We thus represent the probability of a single particle binding to location *x* on the column with the exponential probability distribution, with λ being the average distance a particle travels in solution before being absorbed into the column wall.

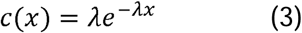

λ is directly dependent on the speed of the mobile phase (µ) and inversely proportional to the diffusion coefficient (*D*) and relative affinity for the column over the solute.

For a single particle, this event does not happen a single time, but many times over the course of column loading. To represent this for *n* binding/unbinding events, we can sum *n* exponential functions together, generating an Erlang distribution^24^.

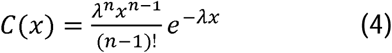

At large *n*, the erlang distribution will converge to a normal distribution^24^ with mean *n*λ and variance *n*λ^2^.

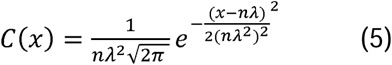

This is the probability distribution for the location of a single particle along the column. To represent the probability distribution of *M* particles, we can multiply the distribution by *M*.

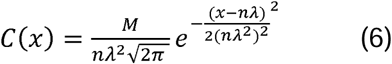

To convert this distribution to an arrival time domain, we divide the distance *x* from the column by the flow rate *µ*.

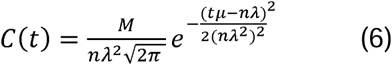

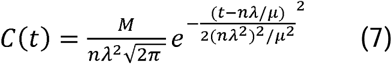

To simplify the expression, we will define two new variables *b* = *nλ*^2^/*µ* and *c* = *nλ*/*µ* These variables are the spatial mean and variance from equation 5 converted to the arrival time domain by dividing by the flow rate *µ*. This transformation yields the following equation:

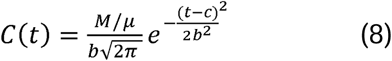

This is a Gaussian distribution, which is supported in the chromatography literature as the canonical distribution for peaks in isotonic elution conditions^8,9^. However, when performing elution over a gradient of solvents, the relative affinity of the analyte for the column and mobile phases shifts: encouraging single-step Langmuir kinetics at a critical point on the gradient near the retention time, which results in the exponential decay behavior with no rebinding, which is observed in equation 1. To combine these two effects, we can convolve the two functions.

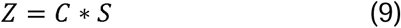

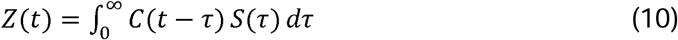

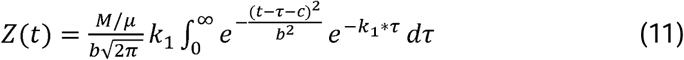

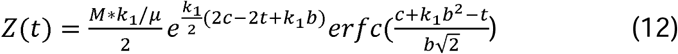

This matches the standard form of the exponential gaussian function, with *erfc* representing the inverse error function 1 − *erf*(*x*), with 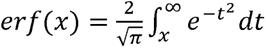. This equation gives us several insights into the factors that influence the shape of the resulting function. The amplitude, or height of the exponential gaussian function is directly proportional to the number of analyte molecules, and inversely proportional to the flow rate. The center *c* of the distribution is dependent on the average travel distance of the particle during column loading and the number of binding/unbinding events, whose dependency has been previously described. The width or *σ* (*b*) is dependent on the same parameters but can also be affected by other minor parameters such as longitudinal diffusion and column inhomogeneities, and thus is not directly proportional to the distribution center. Finally, *k*_1_ is the unbinding coefficient of the analyte from the column and represents the gamma variable, γ, of the exponential Gaussian. This determines how large the tails of the function are. Although our model makes several simplifying assumptions, such as a constant λ during column loading and no longitudinal diffusion, it provides a sound biophysical justification for use of the exponential gaussian distribution, which has been utilized in previous chromatography studies^8,20^. To verify this analytical equation, we conducted a Monte-Carlo simulation recapitulating the assumptions of a period of unbinding/rebinding to the column followed by a kinetic phase in which the analyte has strong affinity for the solvent. This simulation fit an exponential Gaussian equation almost exactly (Figure 2B, red line on blue histogram), further supporting the exponential Gaussian as a good distribution to model HPLC peaks.

#### Empirical Solution

To empirically test our theoretical exponential Gaussian distribution on real data, we injected single, pure fatty acid analytes (linoleic acid, arachidonic acid, and docosahexaenoic acid) onto a C18 column, at individual concentrations of either 0.5mg/mL or 1 mg/mL. Analytes were eluted from the column on a 3:1 methanol water to acetonitrile gradient (see methods) adapted from^25^. We then used the Python package lmfit^26^ to fit one of four functions commonly used in chromatography to each of the fatty acid peaks. A representative chromatograph of linoleic acid is shown to be fit to i). a Gaussian distribution^13^, ii). an exponential Gaussian distribution, iii). a Voigt distribution^27^ and iv). a skewed Gaussian distribution^14^ (Figure 2C). The goodness of fit was calculated using the Bayesian Information Criteria (BIC), which scores models both based on its residual function and the number of parameters. The skew and exponential Gaussian distributions both have an additional parameter as compared to the Gaussian and Voigt functions, making this comparison necessary. The exponential Gaussian distribution had by far the lowest BIC for both concentrations of analytes (Figure 2D).

### PeakClimber software package to rapidly and accurately quantifies chromatography peak areas

We created PeakClimber, an algorithm and python package that identifies and quantifies individual peaks on a chromatographic trace by fitting a sum of exponential Gaussian functions to the HPLC trace (Figure 3).

**Figure 3:**
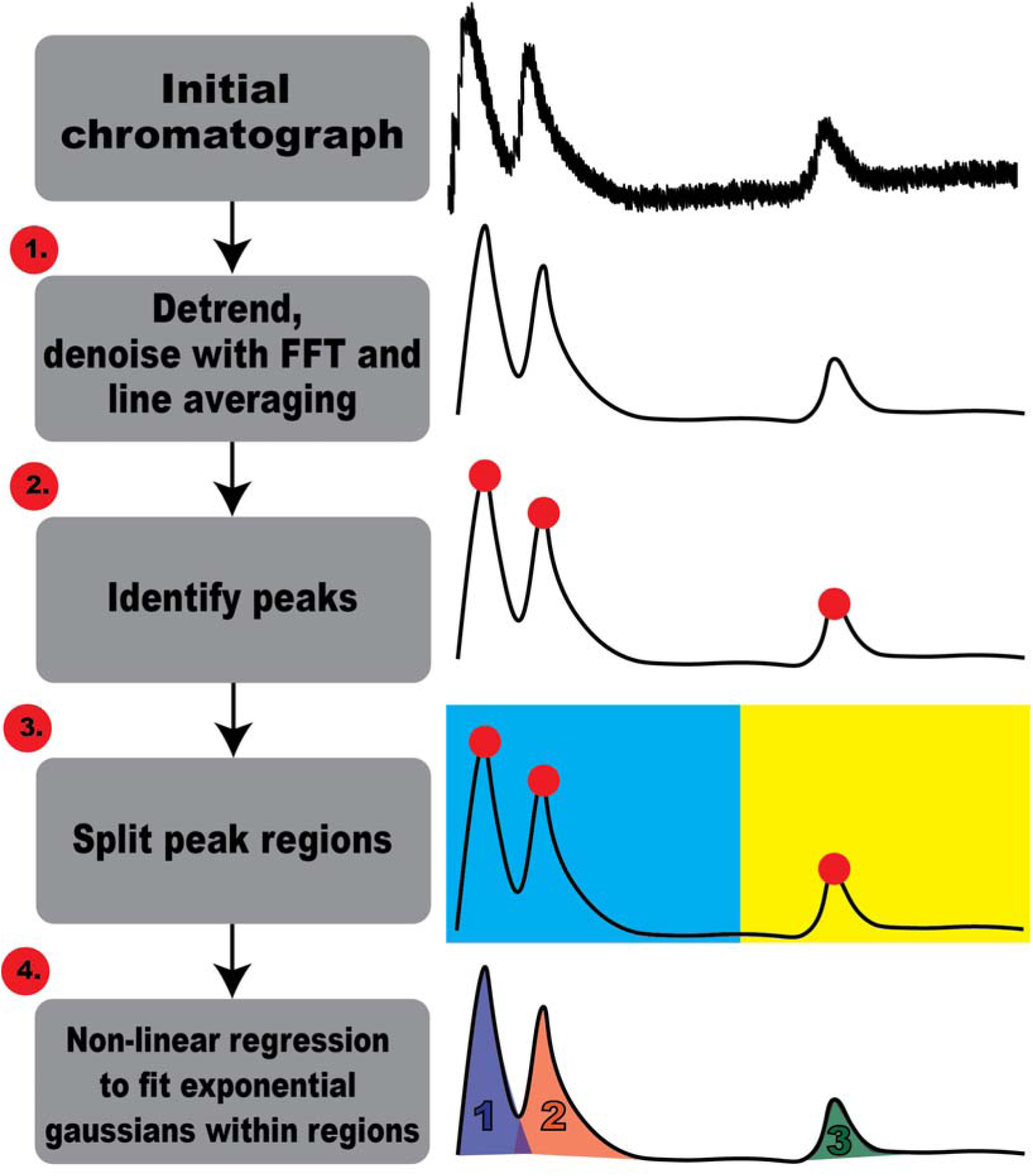
The PeakClimber workflow. (1) PeakClimber first denoises (FFT) and detrends chromatography data before (2) identifying peaks using prominence cutoffs. (3) To decrease runtime, peaks are split into regions based on intersections of the trace with the x-axis. (4) Within each region, peaks are fit to an exponential Gaussian distribution.

Taking a text file of the trace as input, PeakClimber first denoises and detrends the data. Denoising is accomplished using a low pass FFT filter^28^, as well as time-averaging convolution. Detrending is accomplished with a high-pass Whitaker baseline subtraction algorithm that was developed for chromatography, called the peaked signal’s asymmetric least squares algorithm (psalsa)^29^. Exact parameters for these detrending algorithms are input by the user. We chose our default values by fitting single peaks of real HPLC data (Figure 3-1). Peaks are then identified on the denoised data using scipy’s peak finding algorithm, relying on prominence cutoffs to determine if peaks are real or noise^30,31^. The prominence cutoff is also user defined in PeakClimber. In this paper, we use a value of 0.05, meaning peaks must be 5% above the contour trough of surrounding peaks to be analyzed. Additional peaks that form shoulders on more prominent peaks can be optionally identified by identifying local minima and maxima in the first derivative of the HPLC trace that are close to 0 (Figure 3-2).

For each identified peak, an exponential Gaussian function is fit using lmfit^26^ with default parameters of the identified peak center, the identified peak height, a sigma of 0.1 minutes, and an exponential decay parameter of 2. Boundaries between discrete peak regions are set where the background-subtracted trace hits zero (Figure 3-3). The discrete peak regions of the graph are effectively independent of each other, meaning fits can be performed independently on each region without loss of accuracy. Each group of Gaussians is fit to the trace in the appropriate region using a non-linear regression to minimize the least-squared distance between the generated sum of functions and the underlying trace (Figure 3-4). The algorithm recombines the fits for the different windows and returns a summary graph of the resulting fit, overlaid with individual peaks, as well as a table with peak number, runtime, and peak area.

**Figure 4:**
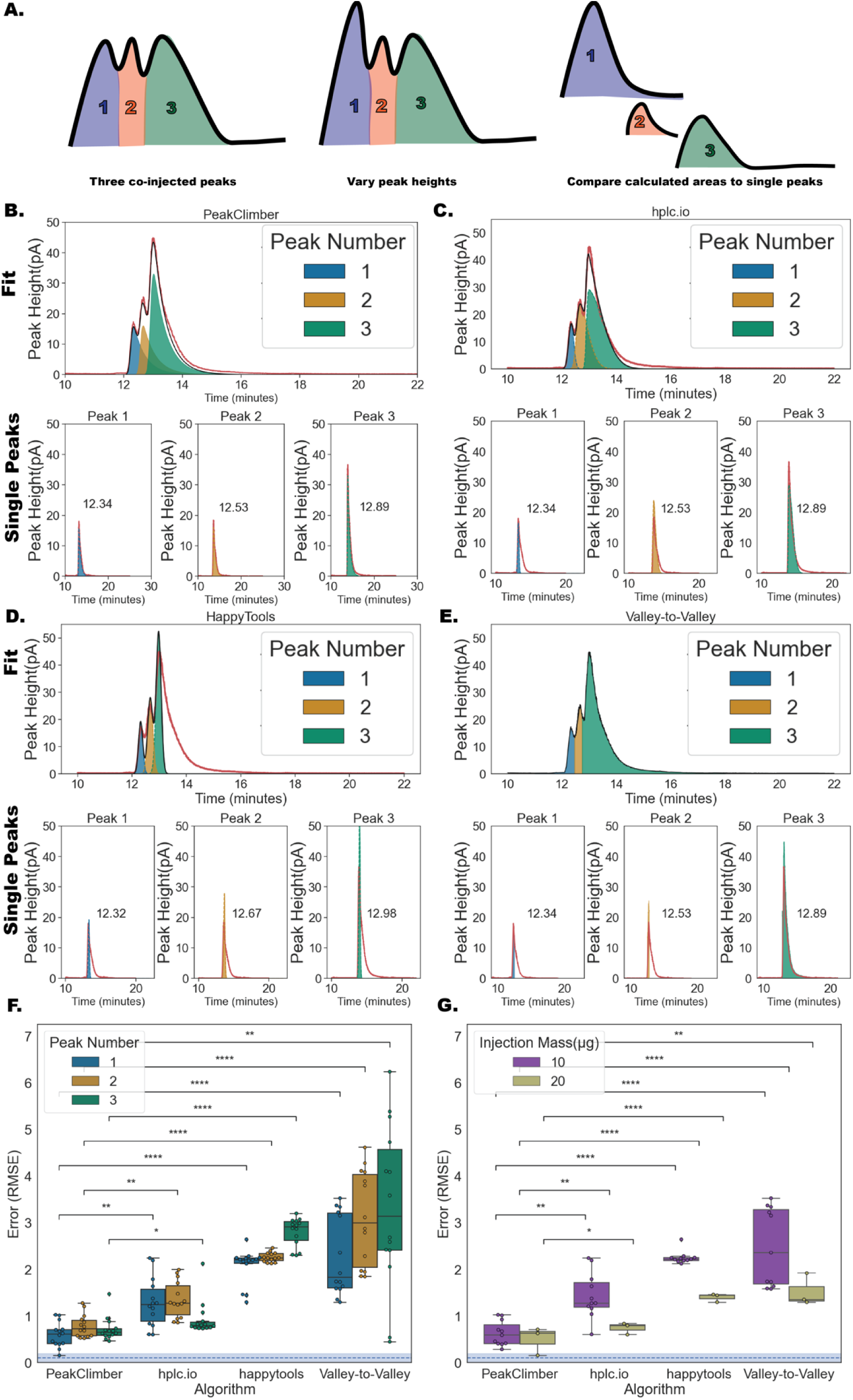
PeakClimber is more accurate and precise than Industry Software. (**A**). Schematic of the experiment performed in this figure. Three overlapping fatty-acid peaks were injected at either a ratio of 1:1:1, 1:2:1, 2:1:1, or 1:1:2. The calculated areas (using the algorithms listed below) were compared to the real injected areas of the individual peaks. (**B**) PeakClimber fit to a chromatograph of a coinjection of C18:1, C20:4 and C22:6 mixed in a 1:2:1 ratio. Red trace: raw data, black trace: predicted sum of peak areas, blue, orange, green shaded regions: predicted individual peak areas. (**C-E**) Fits to the same data in **B** by (**C**) hplc.io, (**D**) the valley-to-valley algorithm (Thermofisher and Waters), and (**E**) HappyTools. Bottom subpanels show the fits to each individual peak. (**F**) Quantification of error rates by RMSE of pooled coinjections of C18:1, C20:4 and C22:6 depending on peak position by above algorithms. Blue dotted line represents minimum RMSE error obtained from the single-peak fits. (**G**) Quantification of error rates for peak 1 alone comparing 10 µg and 20 µg injections. (Kruskal-Wallis Test with Bonferroni correction for subpanels **F** and **G**, n=12, 3 biological replicates each with 4 experimental replicates, *: p<5e-02**: p<1e-02, ***: p <1e-03, ****: p < 1e-04).

### Comparison of PeakClimber to other common HPLC algorithms

To test the utility of PeakClimber, we compared its performance to publicly available software. To generate a test dataset with known standards, we injected three fatty acids with overlapping retention times: docosahexaenoic acid (12.3 minutes), arachidonic acid (12.5 minutes) and linoleic acid (12.9 minutes). We ran the analytes at concentrations of either 0.5 or 1 mg/mL. Thus, in each injection, the analytes were either or equal concentration or one analyte was double the concentration of the other two (Figure 4A). This was done to test the dynamic range of PeakClimber compared to other algorithms. The raw HPLC trace was then smoothed and normalized before fitting the three peaks by one of four algorithms. PeakClimber is the algorithm described in this paper (Figure 4B). hplc.io^14^ is a python-based, chromatographic fitting software that uses skewed Gaussian functions as representative of single peaks (Figure 4C). Happytools is a free, standalone software package that uses Gaussian functions to fit single peaks^13^ (Figure 4D). Finally, valley-to-valley is an abstraction of algorithms^5,6^ used by common HPLC-software such as Thermofisher’s Chromeleon, Agilent’s OpenLab CDS, or Water’s MassLynx that integrates the area under the curve of the trace between “valleys”, the lowest points between two identified peaks (Figure 4E). Fits (black line in Figure 4B-E) were performed on the entire trace (red line in Figure 4B-E). Error comparisons are reported for each individual peak for each of the three analytes (lower panel Figure 4B-E; analytes are DHA, ARA, and LA from left to right) using root mean-squared error (RMSE). Fit peaks were recentered on the canonical single analyte peaks because run times shifted to later elution times in the co-injections. PeakClimber outperformed all other software regardless of peak position (Figure 4F). Particularly for the first peak in the co-injection, PeakClimber has lower error than the other algorithms due to the correct fitting of the tail of the peaks (Figure 4G). PeakClimber also performed better for the second and third peaks (Supplemental Figure 1 A & B). When error is calculated through percent error of the peak area, rather than RMSE, this pattern still holds (Supplement Figure 1 C-F).

### Testing the Limits of Peak Climber

All algorithms, including Peak Climber, have reduced accuracy for groups of peaks under three separate circumstances: high signal-to-noise ratios, small distance between peaks, and uneven ratio between small and large peaks. To test these bounds specifically for PeakClimber, we computationally created traces of partially overlapping peaks using the real fatty acid traces that we generated in Figure 2 with different levels of noise added on top of the trace. The first and second peak overlap, while the third peak is functionally independent, serving as a negative control (Supplemental Figure 2). To test the fitting ability of the algorithm, rather than peak finding, which is already well-tested in other works^30^, we provided the peak locations to each algorithm. The % error for each of these cases is shown in the respective subpanels of Figure 5.

**Figure 5:**
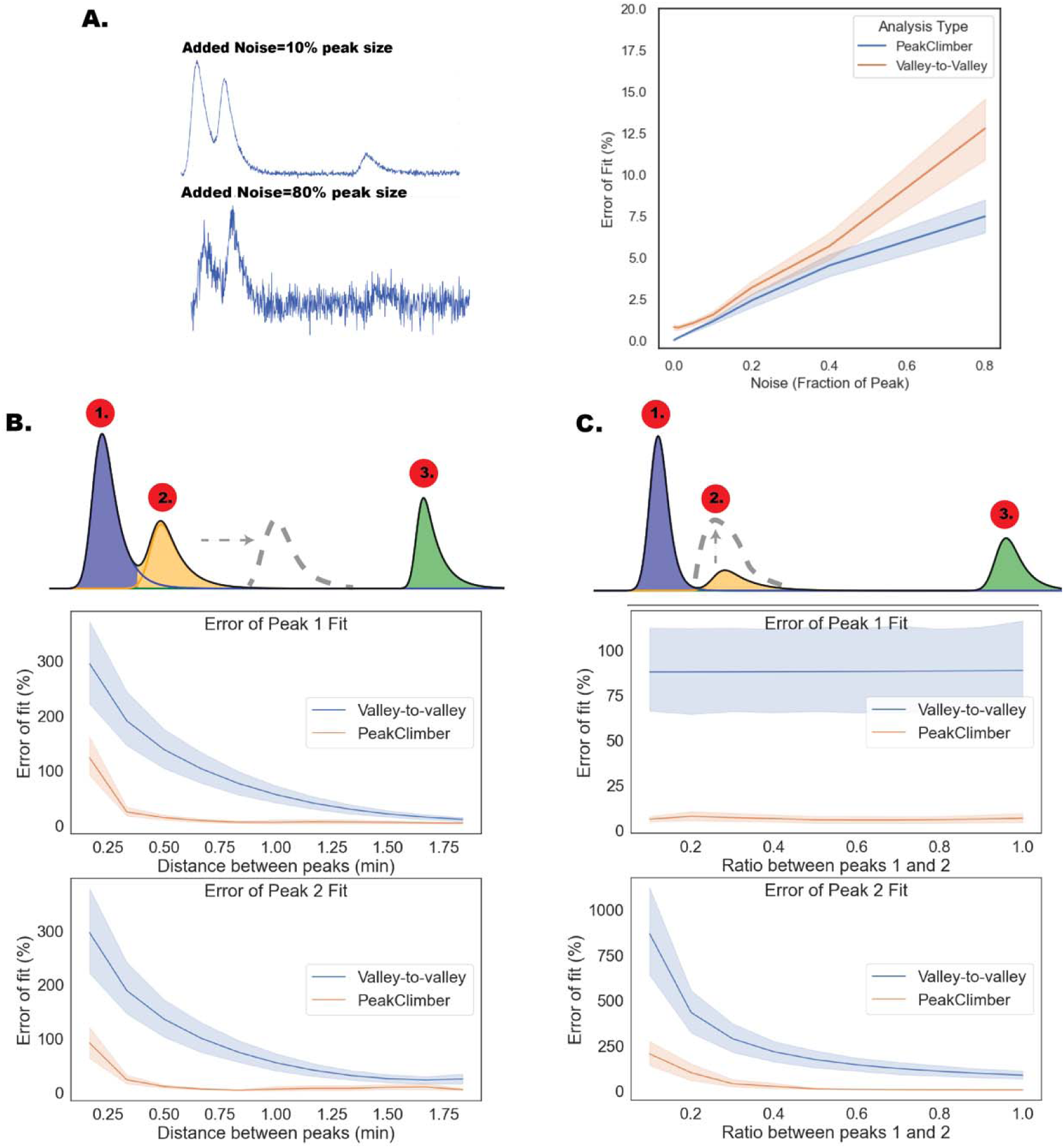
PeakClimber is more robust to noise and peak overlap than industry software. (**A**) Schematic: Gaussian noise as a fraction of the peak amplitude is added to a single synthetically generated peak with amplitude between 1 and 2, a between 0.1 and 0.2, γ between 2.9 and 3 and center at 1. This noise is detrended and removed using the PeakClimber algorithm and then the resulting peak area is either found by fitting (PeakClimber) or integration with the valley-to-valley method. The calculated area is compared to the known area. (**B**) Three analyte curves are superimposed and shifted 0.1-2 min (peak 2) or 10 min later (peak 3). Curves are generated from real traces of arachidonic acid, docosahexaenoic acid, and linoleic acid (**C**). Using the same parameters for exponential Gaussians as in **A**, but with a fixed distance of 0.75 minutes between peaks 1 and 2, and 10 minutes between peaks 1 and 3, the ratio between peak 1 and 2 was varied between 0.1 and 1 (n=24 [4 experimental replicates of each of the 3 fatty-acids at 2 concentrations]).

For noise on single peaks, PeakClimber outperforms manual integration for added noise with an amplitude that is 0.3 times or greater than the true peak size (Figure 5A). This is likely because PeakClimber better captures the shape of the underlying peak. For the distance between peaks, PeakClimber accuracy begins to drop off when the distance between peaks is less than 0.25 minutes. The valley-to-valley method is similarly sensitive to peak overlap only at a threshold distance of 1.5 minutes (Figure 5B). For the ratio between peaks, we held the peaks a fixed distance of 1.5 minutes apart. Varying the said ratio between the first and second peaks did not change the error rate for the larger first peak although PeakClimber outperformed valley-to-valley at every peak ratio. For the second peak, both algorithms have large error rates at ratios below 10:1 large peak:small peak. However, PeakClimber’s error drops rapidly to 0 by a ratio of 4:1, whereas the valley-to-valley method not only drops in error more slowly, but also converges to a steady error rate of about 85% (Figure 5C). This error rate is the lower bound for the valley-to-valley method for peaks with this interpeak distance (Figure 5B).

### Uniqueness of PeakClimber Solution

PeakClimber identifies peak areas by fitting exponential gaussian functions to the underlying chromatography trace using non-linear regression^26^. We can define the error as the sum of residuals between the *y_i_* and the sum of *n* exponential gaussian functions *f_n_*(*x_i_*)

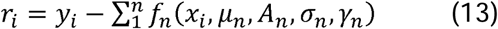

With *µ_n_, A_n_, λ_n_, γ_n_* being the center, amplitude, width, and decay function of each gaussian respectively. This residual function will have a single solution if the second derivatives of the function r are all positive, in other words, if the function is convex. When the shape is of y is equivalent to the sum of exponential gaussians, this function will simplify to 0, which is trivially convex, meaning there is only a single solution.

Additionally, we can empirically restrict the sample space of parameters by observing real behavior of single HPLC peaks. For example, peak centers do not vary from their locations in identified traces, meaning that we can effectively reduce the parameter space down to 3 parameters for each exponential gaussian. Kinetics and diffusion-to-flow ratios also place biophysical limits on the upper and lower bounds for the γ (tail, *k*_1_ in equation eq. 11), and *σ* (width, *b* in eq. 11) parameters. In this reduced parameter space, we find a single optimum for two overlapping exponential Gaussians fit to a region of the lipid profile of *D. melanogaster* thought to contain only two peaks. Since the space is mapped by 6 parameters (not including the fixed centers), we used dimension reduction to visualize the result as a PCA, which shows only a single minimum of the residual *χ*^2^ function (Supplementary Table 1).

### PeakClimber can be used to accurately quantify lipid differences between biological samples

To test the utility of PeakClimber on real biological data, we raised female *Drosophila melanogaster* from the larval stage on two different microbial conditions (germ-free or conventionally reared) on a standard diet. We then performed a lipid extraction and then ran the isolated lipids on the HPLC, using a two-step gradient (first methanol:water to acetonitrile, then acetonitrile to isopropanol) to separate lipid species by polarity and size, as adapted from^25^. Significant differences are observed by eye between germ-free and colonized animals (Figure 6A), especially in the triglyceride region running from 60 to 70 minutes (Figure 6A, inset). Individual peaks were quantified using either the PeakClimber (Figure 6B, left panel) or Thermofisher Chromleon (Figure 6B, right panel). The two algorithms identified the same peaks but produced differences in the magnitude and statistical significance between colonized and germ-free animals (Figure 6C). Chromeleon identifies all peaks in this region as significantly different between samples, whereas PeakClimber only identifies some of these peaks as differentially present. This is not due to sample variance: PeakClimber and Chromeleon both capture biological sample variance equally. This discrepancy is likely because the tail of the first peak contributes to the area counted as the second peak by Chromeleon, causing a false positive when the area is counted this way. This does not occur with PeakClimber, which is able to deconvolve the tail of the first peak from the rest of the second peak. This suggests that PeakClimber has more utility in identifying real differential peaks as compared to standard industry software.

**Figure 6:**
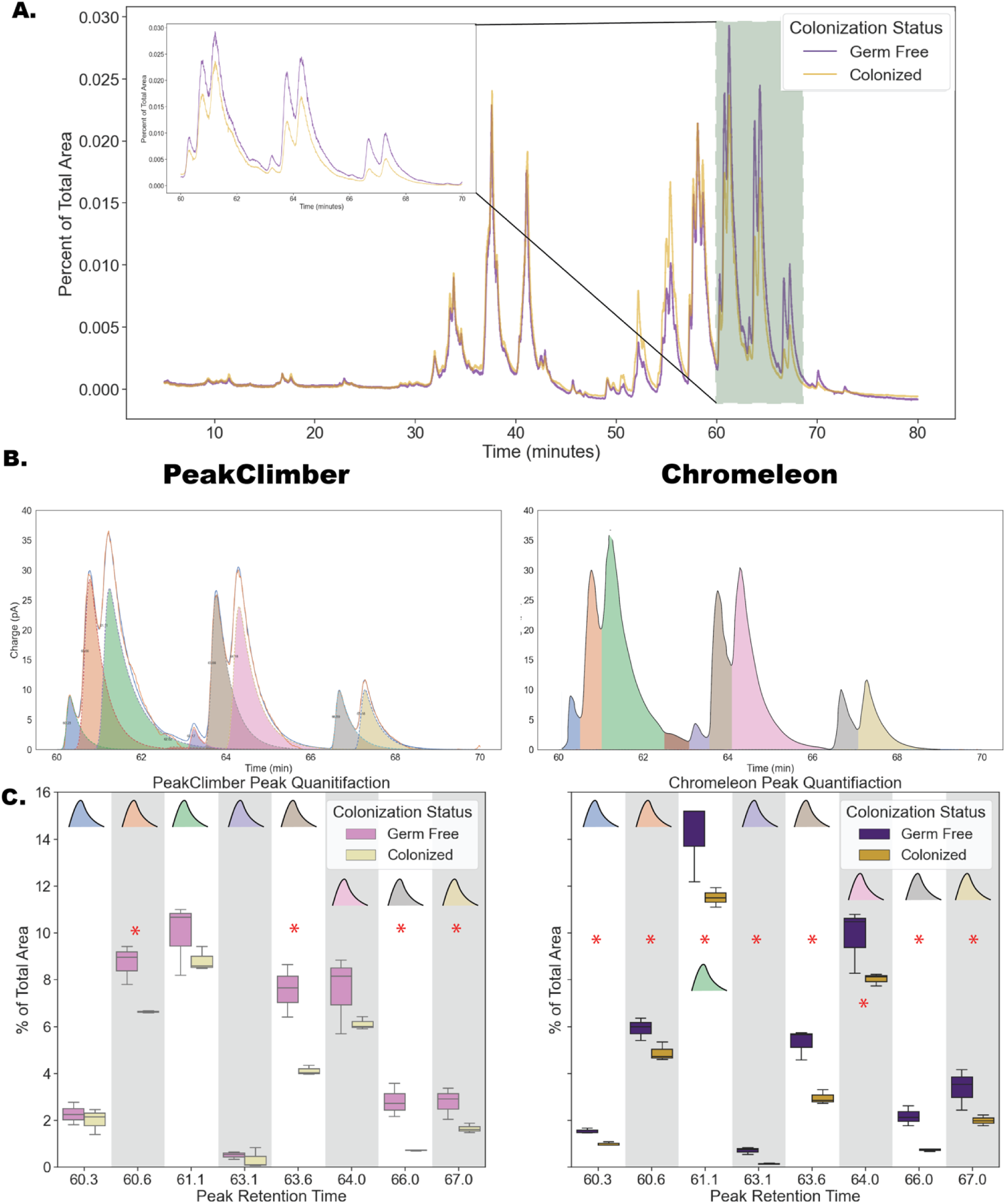
PeakClimber more accurately quantifies biological differences between germ-free and colonized flies. **(A)** A lipid profile for germ free (purple) or colonized (yellow) female fruit flies normalized to total chromatograph area. Highlighted regions in green are shown in higher resolution in the inset are quantified below in (**B**). Algorithmic fit of the highlighted regions above using PeakClimber (left) and Thermofisher Chromeleon (right) on the germ-free trace. **(C).** Quantitation of peak areas for selected peaks in the highlighted region (Krukskal-Wallis rank-sum: *: p< 0.05 n=3, 8 flies per sample).

To identify the lipids contained in these peaks, we first performed a lipidomic-mass spec analysis of whole male and female flies to establish a dataset for canonical fly lipid compounds. Then, we isolated the 8 sample peaks identified in Figure 6B and ran them through a LC-MS system to determine their identities. We used the lipidomic data to verify the LC-MS results from individual peaks. Many m/z numbers from the individual peak analysis were not found in lipidomic-mass spec dataset so we also considered compounds that were more saturated, which would increase the predicted m/z by 1 for each additional hydrogen. Individual peaks were dried down in a vacuum centrifuge overnight, oxidation and subsequent increase in m/z values, could have occurred. We found that except for the first peak that ran at 60.1 minutes, the remaining 7 peaks were triglycerides (Table 1). These peaks were relatively rich in medium-chain triglycerides, which in agreement with other literature on *Drosophila* lipids^32^. Additionally, the specific elution time of these triglyceride peaks nicely agrees with prior HPLC data of zebrafish lipid extracts that were also subject to mass spec confirmation^25^.

**Table 1:** Mass-spec analysis and identification of *Drosophila* lipids eluted from the triglyceride region of the total lipid chromatograph. The retention time (column 1), measured m/z value (column 2), predicted unsaturated m/z value (column 3), and compound identity (column 4), and PeakClimber significance (Krukskal-Wallis rank-sum: *: p< 0.05 n=3, 8 flies per sample) for each peak in the 60-70 minute region of the fly lipid profile (TG=Triglyceride, PC=phosphatidylcholine).

**Figure 6: PeakClimber more accurately quantifies biological differences between germ-free**
| Retention Time | Measured m/z | Predicted unsaturated m/z | Compound ID | Significant |
| --- | --- | --- | --- | --- |
| 60.3 | 654.27/846.67 | 654.56/834.6 (-12 H) | TG(36:1)+NH<sub>4</sub>, PC (40:6)+H | No |
| 60.6 | 820.66 | 818.72 (-2 H) | TG(16:1,16:1,16:1)+NH<sub>4</sub> | Yes |
| 61.1 | 794.65 | 790.69 (-4 H) | TG(14:1,16:1,16:1)+NH<sub>4</sub> | No |
| 63.1 | 768.64 | 768.7 | TG(14:0,14:0,16:0)+NH<sub>4</sub> | No |
| 63.6 | 848.69 | 846.75 (-2 H) | TG(18:3,16:0,16:0)+H | Yes |
| 64.0 | 822.68 | 820.73 (-2 H) | TG(16:0,16:1,16:1)+NH<sub>4</sub> | No |
| 66.0 | 876.72 | 869.75 (-7 H) | TG(21:4,16:0,16:0)+NH<sub>4</sub> | Yes |
| 67.0 | 850.70 | 848.77 (-2 H) | TG(14:0,14:0,18:2)+NH<sub>4</sub> | Yes |

Three out of the four significantly enriched peaks in germ-free animals contained long-chain polyunsaturated fatty acids (63.6,66,67 minutes). None of the non-significant peaks contained any polyunsaturated fatty acid tails, perhaps suggesting that colonized animals more readily metabolize these fatty-acids, or that they are preferentially absorbed by microbes, and are thus lost through feces.

## Discussion

In this paper we have shown three principal findings. First, the exponential Gaussian function is a good fit for HPLC peaks. We showed this both analytically, computationally with Monte-Carlo simulations, and empirically by calculating the error of the fit for various common distributions used in chromatography to fit single analyte peaks. Many previous works from the 1970s and 1980s attempt to analytically work out these solutions, and their models, also approximated an exponential Gaussian distribution ^8,12,15,16,22,33,34^. HPLC peaks often do not represent single compounds, but groups of compounds. This means that a single peak is often a sum of individual compounds, all with behavior as described in **Figure 2**. Due to the central limit theorem, this would suggest that the chromatographic traces that we observe should have more of a Gaussian character, but this is not what we observe empirically.

Second, we demonstrated the effectiveness of PeakClimber as compared to other commercially and freely available software tools to quantitatively analyze chromatography data with overlapping peaks. This is due to the ability of our algorithm to capture the tail region of the first peak in a group of peaks, which prevents undercounting and reduces distortion by larger surrounding peaks. We also show that, given biophysical assumptions that limit the parameter space, there is only a single best fit solution for the underlying trace. This is vital for accurately quantifying peaks.

The package and documentation for PeakClimber are freely available on GitHub with an easy-to-use graphic user interface (GUI). One limitation of the test data used to validate PeakClimber is that it was only used to test HPLC data from lipid chromatography. Theoretically other biomolecules should have the same kinetic and diffusive behaviors as lipids, and many chromatographic traces present in the literature show single peaks that appear to be similar to exponentially modified Gaussian functions^7,8,35–38^. However, adapting our algorithm to additionally deal with anti-Langmuir fronted peaks^39,40^ could be a promising next step.

Third, we demonstrated the utility of our algorithm for the analysis of biological data. While mass-spectrometry will always be the gold standard for metabolic analysis^41^, HPLC represents a lower-cost medium-throughput option than mass-spec^42–44^. Consider an experiment similar to one that we set up with multiple replicates of different dietary, genetic, or microbial conditions. Rather than analyze each replicate by mass spectrometry, one replicate from each group could be run through mass-spec, and the rest on HPLC, where relative changes in the compounds identified by MS could be much more accurately quantified with PeakClimber. The recognition of HPLC as this medium-throughput bridge between MS and high-throughput methods such as colorimetric kits could be one reason for the recent interest in development of tools to better analyze this type of data^2,41,43,44^.

The reduction in triglycerides containing long-chain fatty acids polyunsaturated fatty acids in flies colonized with *Lactobacillus* and *Acetobacter* is an additional interesting finding from this work. Previous work in mice^45,46^ has shown that various *Lactobacillus* species can protect against obesity by acting as a sponge for fatty acids, and then being excreted in the feces. These results also agree with work in the fly that shows that colonization can reduce triglyceride accumulation^47,48^. Why these bacteria reduce the presence of polyunsaturated fats in particular is unclear, but could be due to composition of Lactobacillus membranes, which are largely composed of unsaturated fatty acids^49^.

Although we did not observe this in our dataset, neighboring peaks in HPLC are often composed of extremely similar compounds that are part of biochemical pathways such as fatty acid elongation or conversion between different phospholipid compounds^50,51^. PeakClimber could be used to find the precise step in these pathways that is affected by the genetic mutation, diet, or colonization condition of interest. This method could provide an additional advantage over alternative methods such as transcriptomic or proteomic analysis due to the ability to measure actual metabolite levels rather proxies of RNA or protein levels, the activity of which can both be affected by downstream processing such as translation (in the case of RNA), or post-translational modifications and confirmational changes (in the case of protein).

### Limitations and comparison to other algorithms

The exponential Gaussian function will not perfectly fit some chromatography peaks, as compounds that run with anti-Langmuir kinetics will have peak fronting^15,52,53^, which will not fit an exponential Gaussian distribution. Peak fronting can also occur when the peak has been overloaded with analyte. Additionally, our mathematical model makes several simplifying assumptions about the geometry and flow rate of common HPLC systems. Based on complexities of experimental conditions that influence the quality of the data, more complex analysis programs could be needed.

However, despite these limitations, we want to highlight the performance advantages of PeakClimber compared to other available software. Prior to PeakClimber, there was a common sentiment that overlapping peaks could only be analyzed qualitatively. Here with PeakClimber we show that we can in fact extract highly quantitative data from HPLC traces that contain overlapping peaks. Even compared to other software that attempts to tackle this problem in a similar manner, PeakClimber much more accurately quantifies areas of overlapping peaks, due to its ability to consider long peak tails. For the analysis of biological data that contain many overlapping and non-overloaded peaks, we believe that PeakClimber will prove to be invaluable.

## Materials and Methods

### Monte-Carlo simulations

#### Exponential decay simulation

A column 1000 units long with 100 analyte particles at position 1 bound to the column is instantiated. At each time step the analyte has a 5% chance of unbinding from the column (representing a k value of 0.05). Once unbound the particle arrives at the detector a fixed time later, in this case 900 time-steps. The simulation was performed 10000 times and results were pooled.

#### Multi-step reaction simulation

A column 1000 units long with 100 analyte particles at position 1 is instantiated. Particles are allowed to bind to the column with a probability of 0.5 and unbind with the same probability for 100 steps. When not bound to the column, particles move at the flow rate (1 binding site per step). After 100 steps, the probability of unbinding is reduced to 0.05, and the probability of rebinding is reduced to 0. Once unbound, these particles arrive at the detector a fixed time later, in this case 900 time-steps. The simulation was performed 10000 times and results were pooled.

### Fatty-acid chromatography

Fatty acid aliquots were obtained from Cayman Chemicals: linoleic acid (LA) (CAS 60-33-3), arachidonic acid (ARA) (CAS 506-32-1), and docosahexaenoic acid (DHA) (CAS 6217-54-5). The fatty acids were suspended in HPLC-grade isopropanol in stock concentrations of 10mg/mL. Aliquots were then further diluted to either 0.5mg/mL or 1 mg/mL as individual analytes or as part of one of the four mixtures analyzed (0.5:0.5:0.5, 1:0.5:0.5, 0.5:1:0.5, 0.5:0.5:1 mg/mL of DHA: ARA: LA respectively. 20 µL of each individual analyte or mixture was injected onto the HPLC system. The components of each sample were separated and detected by an HPLC system using a LPG-3400RS quaternary pump, WPS-3000TRS autosampler (maintained at 20°C), TCC-3000RS column oven (maintained at 40°C), Accucore C18 column (150 × 3.0 mm, 2.6 μm particle size), FLD-3100 fluorescence detector (8 μL flow cell maintained at 45°C), and a Dionex Corona Veo charged aerosol detector (all from Thermo Fisher Scientific). Component peaks were resolved over a 30 min time range in a multistep mobile phase gradient as follows: 0–5 min = 0.8 mL/min in 98% mobile phase A (methanol-water-acetic acid, 750:250:4) and 2% mobile phase B (acetonitrile-acetic acid, 1,000:4); 5–30 min = 0.8–1.0 mL/min, 98–30% A, 2–44% B, and 0–3.3% mobile phase C (2-propanol)^32^. HPLC-grade acetic acid and 2-propanol were purchased from Fisher Scientific and HPLC-grade methanol and acetonitrile were purchased from Sigma-Aldrich.

### Error tolerance simulations

#### Noise simulation

A single exponential Gaussian peak was initialized with the following parameters: amplitude between 1 and 5, γ (skew) between 2.9 and 3, and sigma between 0.1 and 0.2. Noise was added to the peak between 0 and 80% of its amplitude. Peak area was calculated using PeakClimber or manual integration after denoising and compared to the known area of the generated peak.

#### Proximity and ratio simulations

Individual analyte traces of either linoleic acid (LA) (CAS 60-33-3), arachidonic acid (ARA) (CAS 506-32-1), or docosahexanoic acid (DHA) (CAS 6217-54-5) were smoothed and background subtracted as described in the body of the paper. Three copies of the corrected trace were superimposed on top of each other, and the resulting three peaks were computational separated by 0.1 to 2 minutes, or 10 minutes, respectively. The area of each of the three peaks was calculated either using PeakClimber, or the valley-to-valley algorithm, and the compared to the known underlying peak. Error was calculated by dividing the chi-square function of the residual error by the total peak area. For peak ratio, the second peak was held at a constant distance of 0.75, but the relative size of the peak was scaled between 0.1 and 1 of the size of the first peak.

### Fly husbandry

*Drosophila melanogaster* Canton-S flies were initially isolated from long term germ-free stocks kept in lab. The parental generation of flies was inoculated with a 7-species microbiome mixture consisting of *L. plantarum, L. brevis, A. pomorum, A. orientalis, A. cerevisiae, A. sicerae*, and *A. tropocalis* that recapitulates the microbiome found in a wild fruit fly 5 days after eclosion or maintained germ-free. Parental flies were fed a diet consisting of 10% glucose (v/v), 0.42% propionic acid (v/v), 1.2% (w/v) agar, and 5% yeast (w/v). These flies were allowed to lay eggs on their food for 3 days. The resulting offspring were raised until 10 days post eclosion before lipid extractions.

### Lipid extractions and chromatography for *Drosophila melanogaster*

Groups of 8 flies were macerated using a bead beater in 500 µL of lipid extraction buffer (10 mM Tris, 1mM EDTA, 7.8 pH). 400 µL of extract was mixed with 1.5 mL 2:1 chloroform:methanol (with 1 ng/mL of TopFluor cholesterol as an internal standard) and allowed to sit for 10 minutes. Then 500 µL of chloroform followed by 500 µL of extraction buffer was added to the mixture. The mixture was centrifuged at 2300 rcf for 5 minutes and the organic (bottom) phase was harvested. This was evaporated to dryness under vacuum centrifugation and then resuspended in 100 µL of HPLC grade isopropanol.

20 µL of the sample was injected onto the HPLC system as described earlier. Component peaks were resolved over an 80 min time range in a multistep mobile phase gradient as follows: 0–5 min = 0.8 mL/min in 98% mobile phase A (methanol-water-acetic acid, 750:250:4) and 2% mobile phase B (acetonitrile-acetic acid, 1,000:4); 5–35 min = 0.8–1.0 mL/min, 98–30% A, 2–65% B, and 0–5% mobile phase C (2-propanol); 35–45 min = 1.0 mL/min, 30–0% A, 65–95% B, and 5% C; 45–73 min = 1.0 mL/min, 95–60% B and 5–40% C; and 73–80 min = 1.0 mL/min, 60% B, and 40% C. (HPLC-grade acetic acid and 2-propanol were purchased from Fisher Scientific and HPLC-grade methanol and acetonitrile were purchased from Sigma-Aldrich.)

For downstream LC-MS analysis, peaks were collected in the following intervals via fraction collector 60-60.4 min (1), 60.4-60.9 min (2), 60.9-61.5 min (3), 61.5-63 min (4) 63.2-64 min (5),64-66 min (6),66-67 min (7), 67-69 min (8). Peaks were then evaporated to dryness and resuspended in 100 µL methanol and DCM (50/50 v/v) with a concentration of 5mM ammonium acetate in the final solution.

### Liquid Chromatography-High Resolution (Q-TOF) Mass Spectrometry Analysis

Ammonium acetate, methanol, and dichloromethane (DCM) were purchased from Thermo Fisher Scientific Inc. (Waltham, MA). Full scan mass spectral analyses of isolated peaks were conducted using an AB Sciex Quadrupole Time of Flight Mass Spectrometer controlled by Analyst 1.8 (5600 Q-TOF) (Framingham, MA). The mass spectrometer was coupled to a Shimadzu ultrafast liquid chromatographic system (UFLC, Kyoto, Japan), which consisted of a degasser, a quaternary pump, an autosampler, and a temperature-controlled column compartment. Each individual peak fraction (20 µL injection volume) was directly infused into the mass spectrometer’s electrospray ionization (ESI) source chamber through the UFLC autosampler. The mobile phase comprised methanol and DCM (50/50, v/v) spiked with 5 mM ammonium acetate, with a flow rate set to 100 µL/min. ESI parameters were as follows: source gases were set to 20 for Gas 1 and 30 for Gas 2, while the curtain gas was set to 30. The source temperature was set to 250 °C, and the Ion Spray Voltage Floating (ISVF) was set to 5.5 kV. The compound decluttering potential was set to 80. The mass spectrometer operated in TOF high-resolution full scan mode within an m/z scan range of 100 to 1200, with an accumulation time of 0.25 seconds for one minute for each sample run. The high-resolution mass spectrometer was calibrated with manufacturer solvent (PI: 4460131) to maintain mass accuracy.

Canonical lipid peaks were obtained with the following method. 8 frozen *Drosophila melanogaster* females (10 days post eclosion) were resuspended in MTBE (1mL), vortexed and then transferred to an Eppendorf tube. 300 µL of methanol with internal standard was added and samples were shaken for 10 min. 200 µL of water was added to facilitate phase separation.

The extracts were centrifuged at 2,000 rcf for 10 min. The top layer was removed, dried down, and reconstituted in 100 µL of IPA for analysis. Avanti’s deuterated lipid mix, Equisplash, was used as an internal standard. This was spiked into the methanol at 1.5 µg/mL and used for extraction. Analysis was performed using a Thermo Q Exactive Plus coupled to a Waters Acquity H-Class LC. A 100 mm x 2.1 mm, 2.1 µm Waters BEH C18 column was used for separations. The following mobile phases were used: A-60/40 ACN/H20 B-90/10 IPA/ACN; both mobile phases had 10 mM Ammonium Formate and 0.1% Formic Acid.

A flow rate of 0.2 mL/min was used. Starting composition was 32% B, which increased to 40% B at 1 min (held until 1.5 min) then 45% B at 4 minutes. This was increased to 50% B at 5 min, 60% B at 8 min, 70% B at 11 min, and 80% B at 14 min (held until 16 min). At 16 min the composition switched back to starting conditions (32% B) and was held for 4 min to re-equilibrate the column.

### Code availability

All data and Jupyter Notebooks used to generate figures in this manuscript can be found at github.com/ATiredVegan/PeakClimberManuscriptRepository. The PeakClimber package and use instructions can be found at github.com/ATiredVegan/PeakClimber.

## Supporting information

Suppelemental Table 1

Figure catpions for supplement

## Acknowledgements

The authors would like to acknowledge Dr. Brandie Ehrmann of UNC for helping with the generation of the canonical fly lipid profile. The authors would also like to thank Dr. Huiqiao Pan and Dr. McKenna Feltes for testing the PeakClimber GUI. Finally, the authors would like to acknowledge the Farber and Ludington labs for their helpful feedback conceptually and on the actual body of the manuscript.

## Author Contributions

Conceptualization: JTD, SAF, WBL; HPLC: JTD; Mass-Spec: JTD, PD, NH; Fly Husbandry: JTD; Simulations: JTD; Code creation and maintenance: JTD; Figure generation: JTD, WBL; Manuscript drafting: JTD; Manuscript editing: JTD, WBL, SAF.

