## Supplementary material for "PeakClimber: A software tool for the accurate quantification of complex multianalyte HPLC chromatograms using the exponential Gaussian function": Figure catpions for supplement

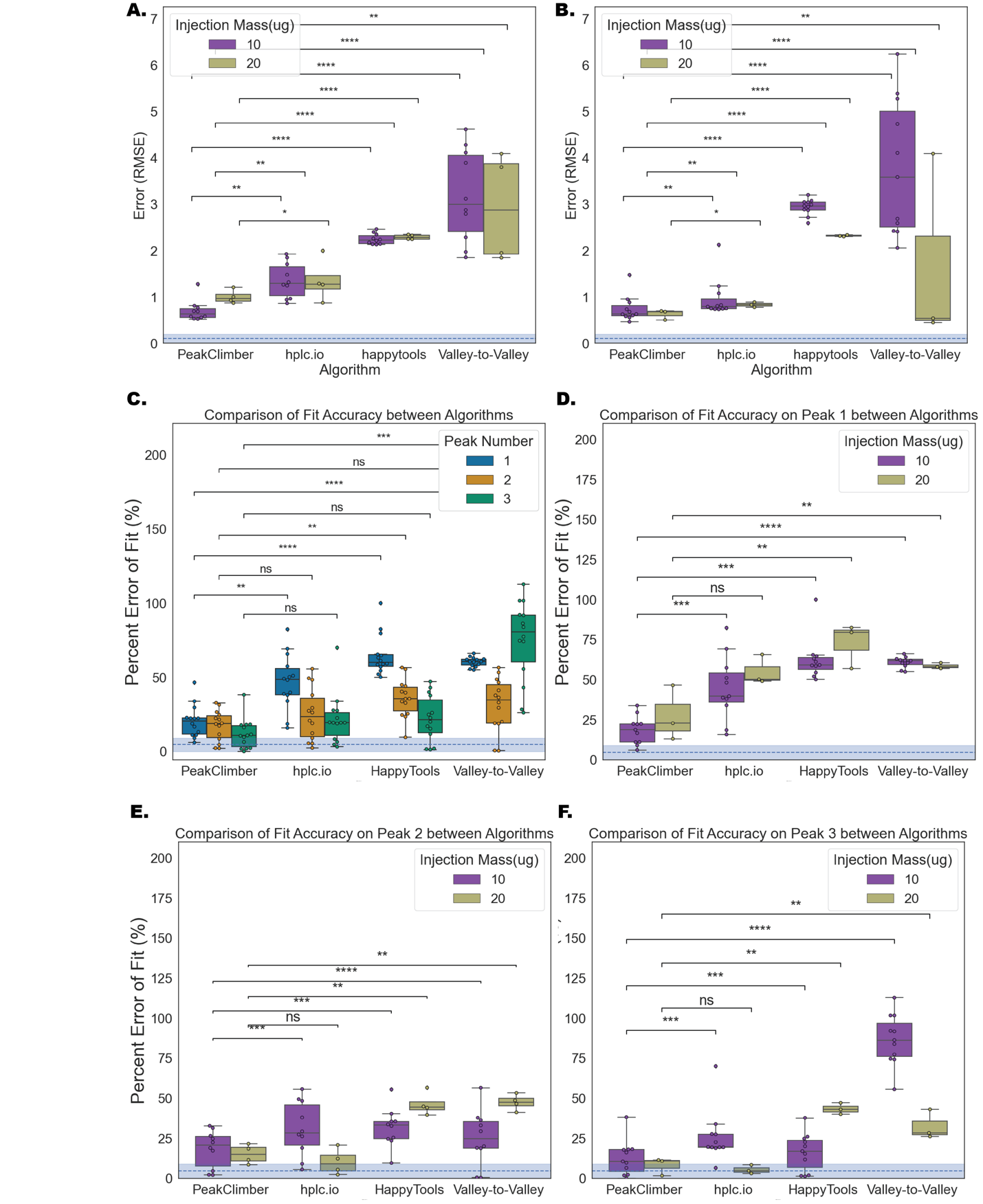

**Supplemental Figure 1: Additional error quantification for PeakClimber.** Quantification of error rates by RMSE of pooled coinjections of C18:1, C20:4 and C22:6 for peak 1 (**A**) and 2 (**B**) error comparing 10 µg and 20 µg injections. (**C**) Quantification of error rates by % error of pooled coinjections of C18:1, C20:4 and C22:6 depending on peak position by above algorithms. Blue dotted line represents minimum % error obtained from the single-peak fits. (**D-F**) Quantification of error rates for peak 1 (**D**), peak 2 (**E**), peak 3 (**F**) alone comparing 10 µg and 20 µg injections. (Kruskal-Wallis Test with Bonferroni correction n=12, 3 biological replicates each with 4 experimental replicates, *: p<5e-02**: p<1e-02, ***: p <1e-03, ****: p < 1e-04).

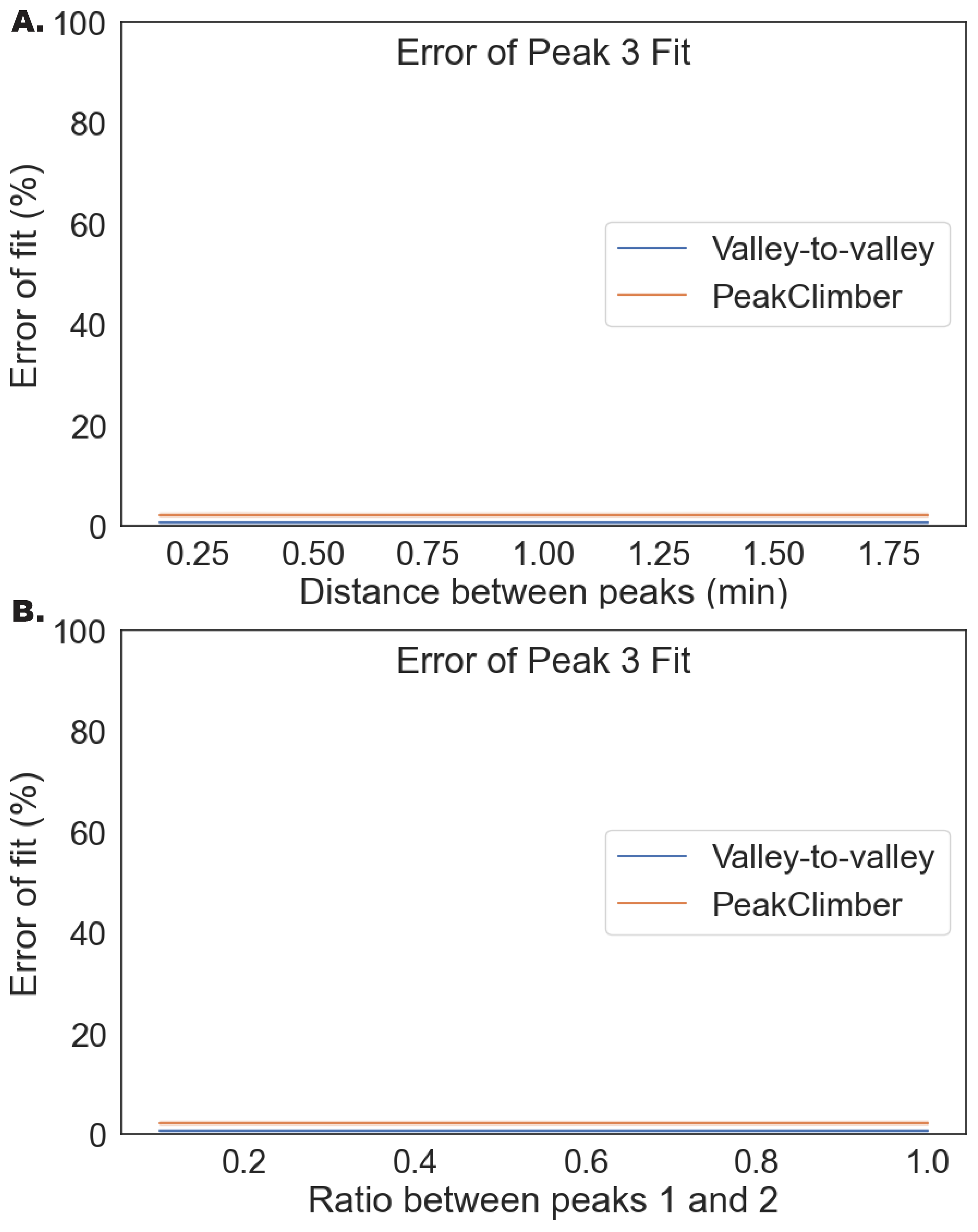

**Supplemental Figure 2: Error tolerance for peak 3**. (**A**). Three analyte curves are superimposed and shifted 0.1-2 min (peak 2) or 10 min later (peak 3). Curves are generated from real traces of arachidonic acid, docosahexaenoic acid, and linoleic acid (**B**). Using the same parameters for exponential Gaussians as in **A**, but with a fixed distance of 0.75 minutes between peaks 1 and 2, and 10 minutes between peaks 1 and 3, the ratio between peak 1 and 2 was varied between 0.1 and 1. In both (**A**) and (**B**) the error rate for only peak 3 is shown. (n=24 [4 experimental replicates of each of the 3 fatty-acids at 2 concentrations]).
