## Supplementary figures and images for "PeakClimber: A software tool for the accurate quantification of complex multianalyte HPLC chromatograms using the exponential Gaussian function"

### Supplemental Figure 1

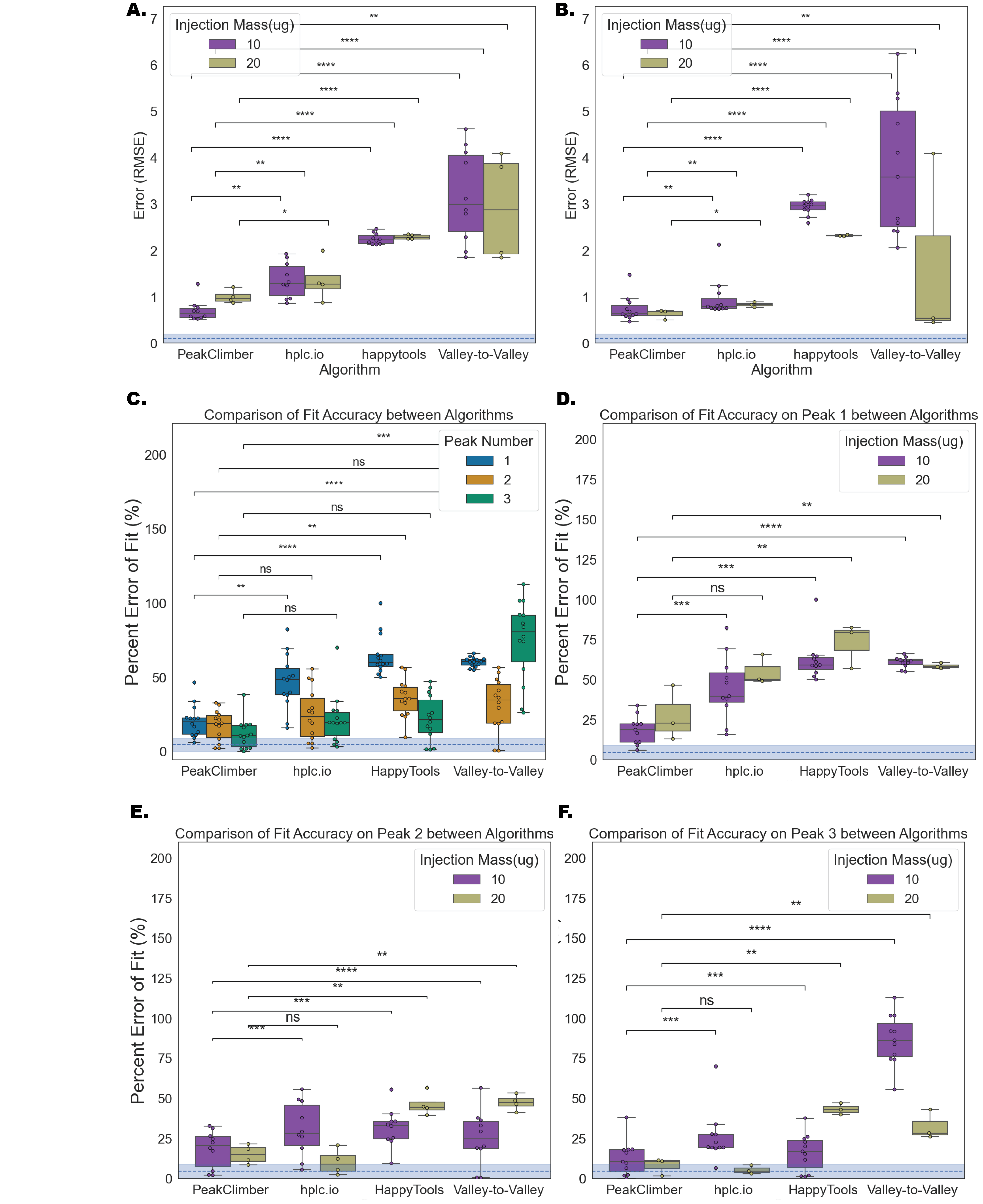

### Supplemental Figure 2

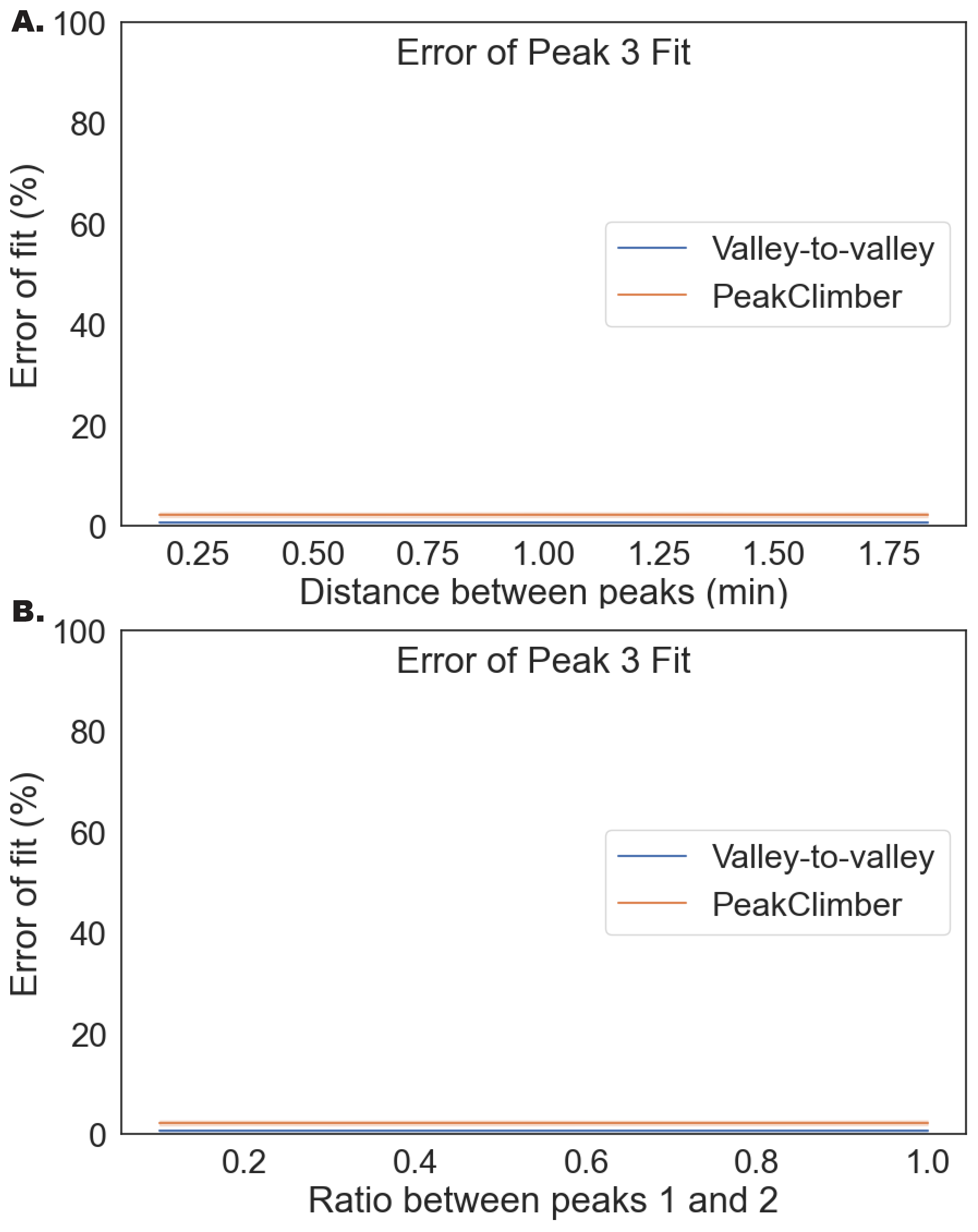
